# Molecular response of the diatom *Coscinodiscus granii* and its co-occurring dictyochophyte during *Lagenisma coscinodisci* parasite infection

**DOI:** 10.1101/2025.10.10.681168

**Authors:** Céline Orvain, Laurie Bertrand, Alice Moussy, Betina M. Porcel, Marine Vallet, Quentin Carradec, Adrien Thurotte

## Abstract

Parasitic interactions play a central role in shaping phytoplankton community dynamics. Diatoms are a major phytoplankton group for which many parasites have been describe including chytrids and oomycetes, yet host defense mechanisms remain poorly studied, limiting our understanding of the factors that constrain or promote infection events in natural environments. Major challenges in investigating diatom-parasite interactions include obtaining cultivable host-parasite pairs, maintain stable co-cultures with synchronized infection stages, and harvesting sufficient biomass for molecular analyses such as transcriptomics and metabolomics.

To address these challenges, we focused on the bloom-forming diatom *Coscinodiscus granii*, a large species (≈200 µm) to allow manual isolation of single cells. This diatom is naturally infected by *Lagenisma coscinodisci*, an abundant oomycete occasionally observed in temperate coastal environments.

We assembled high-quality transcriptomes for both *C. granii* and *L. coscinodisci*, providing an important resource for future molecular studies. Transcriptome analyses revealed a sophisticated effector repertoire in *L. coscinodisci*, including canonical oomycete virulence factors such as Crinklers, RxLR effectors, cystatins, transposon-associated proteins, and components of the RNA interference machinery (Argonaute, Dicer, RdRP), as well as cyclophilins.

In the differential gene expression analyses, *C. granii* exhibited a transcriptional response involving proteases and exosome-related pathways, suggesting a deeply conserved, defense mechanism. In parallel, we analysed the differential expression of the heterotrophic flagellate *Pteridomonas sp*., which consistently co-occurred in culture, and identified a distinct transcriptional profile characterized by the upregulation of motility-related genes, highlighting a striking mobility strategy. Owing to the exceptionally large host size and the availability of both transcriptomic and metabolomic data, this tripartite system provides a unique marine model for exploring oomycete-diatom interactions.

## Introduction

Parasitism is acknowledged as one of the most prevalent ecological strategies among heterotrophic organisms [1]. Protist parasites account for over 50% of the richness and abundance within the heterotrophic pico-nanoplankton (0.8–5 μm) [2]. Parasite organisms often act as parasitoids, killing their unicellular hosts, and can sometimes exert top-down control on microalgal populations, thereby influencing host’s bloom dynamics and termination events [3,4]. In other cases, they are not directly lethal, but may lead to sterility or affect the host’s physical condition and recruitment rates.

Several of the eukaryotic parasites infecting microalgae have been identified in marine plankton. Such is the case of *Amoebophrya*, a wide-ranging and varied group of Syndiniales (also known as marine environmental honeycombs or MALVs [5]) infecting and controlling dinoflagellate populations. Exploring key elements in the interaction with the host throughout their life cycle has shed light on their infection strategies [6–8]. However, underrepresentation of fungal and other eukaryotic parasites in the marine environment (e.g., chytrids, oomycetes, Perkinsozoa) further limits our understanding of their life cycles, host interactions, and ecological roles [9,10].

Diatoms, responsible for approximately 40% of the oceanic export of particulate organic carbon [11] are no exception. They are particularly susceptible to infection by a variety of parasitic eukaryotes, including dinoflagellates, rhizarian, aphelida, and stramenopiles such as hyphochytrids and labyrinthulomycetes [12–14]. Among diatoms, the class Coscinodiscophyceae includes large centric species such as *Coscinodiscus granii*, that commonly form dense, seasonal blooms in coastal environments [15]. These blooms are targeted by various parasitic organisms, including the oomycetes (stramenopiles), a group of heterokont parasites known for their roles in aquatic and terrestrial systems [16]. This is the case of *Pirsonia diadema*, a parasitoid nanoflagellate belonging to the Stramenophila group, which selectively infects *C. granii* by chemosensory stimulation, mechanical attachment and cell wall penetration. These Pirsonia infections can rapidly collapse host populations and potentially influence global biogeochemical cycles [17]. An intriguing feature of this complex interaction is the preference for already infected cells, which increases the potential for subsequent infection [18], as does the influence of environmental factors such as turbulence in the infection.

Another oomycete, *Lagenisma coscinodisci*, has been identified as a specific parasite of *Coscinodiscus granii* species [19]. Unlike most diatom-infecting oomycetes, *L. coscinodisci* develops true hyphae, an elongated filamentous structures that grow within the host and facilitate intracellular colonization and nutrient absorption and has been molecularly characterized as an early-diverging oomycete lineage, providing insights into the early evolution of this group [19,20]. *L. coscinodisci* contributes to *C. granii* bloom decline by massive host lysis and facilitate nutrient recycling within the ecosystem [15]. The *C. granii-L. coscinodisci* system is an emerging model for studying bloom termination. Despite its ecological relevance, there is currently a lack of genomic data for this host–parasite pair, highlighting the need for further molecular and genomic investigations.

In the first molecular insight study of the oomycete *L. coscinodisci*, a combination of co-cultivation experiments and comparative metabolomics approach revealed how the parasite manipulates the metabolome of its diatom host *C. granii* to enhance its infection success [21]. During infection, the oomycete produces specific alkaloids, such as β-carbolines, which accumulate in the reproductive form of the parasite. These compounds inhibit algal cell division, promote plasmolysis, and increase the infection rate, demonstrating a sophisticated hijacking of host metabolic pathways to facilitate the oomycete’s proliferation [21].

In parasitism studies, a substantial limitation to transcriptomics and metabolomics are the presence of asynchronous infection stages in batch cultures [17,22]. In our study, we overcome this obstacle by performing sorted-cell transcriptomics on individually infected *C. granii* cells. To apprehend community chemical sensing, we also exposed *C. granii* healthy cells to infected *C granii* cells in the lytic phase. We assembled the first transcriptomes of C. granii, L. coscinodiscus and an unexpected co-occuring dictyochophyte *Pteridomonas sp*. to explore the molecular response of the diatom host and its associated protists. We conducted a differential expression analysis of the host and its co-occuring Pedinellales (dichtyochophyte), aiming to characterize potential opportunist, escape, or defense responses and their role within this tripartite microbial interaction.

## Material and methods

### Strains and cultivation

Marine diatom *Coscinodiscus granii*, Helg2019-cos20 strain, and the pathogenic oomycete *Lagenisma coscinodisci*, LagC19 strain, were collected from Helgoland, Germany, a location in the North Sea, during the phytoplankton bloom in November 2019. Guillard’s f/2 enrichment medium (G9903, Sigma-Aldrich, Munich, Germany) was prepared by adding 1% of the enrichment solution to 0.22 µm filtered and autoclaved natural seawater (Helgoland, AWI, Bremenhaven, Germany). The cultures were maintained under LED lights at 100 µmol photons m^−2^ s^−1^ 100 of irradiance with a 14:10 day-night cycle and a temperature of 16 °C during the day and 12 °C at night. Microscopy was performed using an inverted microscope (AE30, Motic) for regular checks and sub-cultivation, and a Zeiss Imager 2 (Carl Zeiss) to record microscopic images.

### Infection experiments and sampling

To initiate infection in the diatom host, healthy *C. granii* cells were co-cultivated with infected cells of *L. coscinodisci* by transferring 40 µL of cells in late infection stage (sporangia) into 40 mL of healthy algal cell culture. This involved transferring single diatom cells into a culture with the pathogen, allowing the oomycete to attach and penetrate the algal cells. The infection was maintained by regularly inoculating healthy cultures with aliquots of infected cells every 7 days. The infection progressing in diatom cells was monitored using light microscopy. For the exposition experiment, non-infected *C. granii* were cultivated in parallel to an infected culture using a dual-chamber cultivation system, following established protocol in [21,23].

Due to the asynchronous nature of the infection process, we manually isolated 3 x 100 infected cells at the sporangium stage using an aspiration micro-syringe under the microscope to ensure homogeneity and synchronous samples in the same infection stage, making up the infected algal samples. Infected algal cells in the sporangium stage were selected and isolated when a fully developed hyphae was visible within (Figure 1). The strains used in this study are publicly available and can be obtained free of charge for research purposes upon request to M. Vallet.

**Figure 1.**
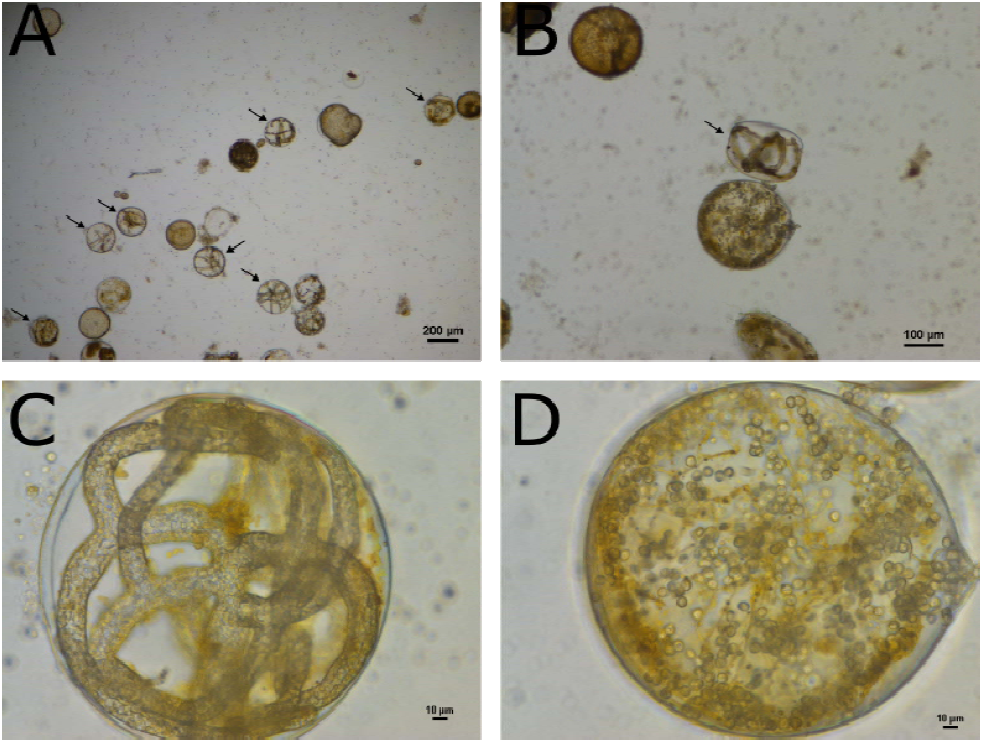
*C. granii* culture observed under bright field microscope. Infected algal cells in the sporangium stage were selected and manually isolated when a fully developed hyphae (C) was visible within (cells with black arrow, A and B). Mature stage cells were not sampled (D).

### Sampling, RNA extraction, cDNA libraries construction and sequencing

Flash-frozen pellets were extracted using RNeasy Plus Universal Mini Kits (Qiagen, Ref 73404). All extracted RNA samples were treated with 6U of TURBO DNase (2 U/µL) (Thermo Fisher Scientific, Ref. AM2238), then purified with RNA Clean and Concentrator-5 kit (Zymo Research, Ref. ZR1016), keeping only large RNA molecules larger than 200 nucleotides for RNAseq library preparation. A total of 30 ng of purified RNA was used to construct Illumina libraries (Illumina Stranded mRNA Prep, Ligation). Briefly, poly(A)+⍰RNAs were selected with oligo(dT) beads, chemically fragmented by divalent cations under high temperature, then converted into single-stranded cDNA using random hexamer priming. A pre-index anchor was ligated, and a PCR amplification step with 18 cycles was conducted to add 10 bp unique dual index adapter sequences (IDT for Illumina RNA UD Indexes, Ligation). All libraries were quantified using Qubit dsDNA HS Assay measurement. A size profile analysis was performed in an Agilent 2100 Bioanalyzer (Agilent Technologies, Santa Clara, CA, USA). Libraries were sequenced in 2 × 150 bp on an Illumina NovaSeq 6000 sequencer (Illumina, San Diego, CA, USA) to obtain 30 million paired-end reads per sample. For infected samples, we sequenced 60 million paired-end reads to ensure sufficient coverage of the additional oomycete. After Illumina sequencing, reads that passed the Illumina quality filters were processed to remove sequencing adaptors and primer sequences, then low-quality nucleotides (*Q* < 20) were discarded from both ends of the reads. Sequences between the second unknown nucleotide (N) and the end of the read were also trimmed. Reads shorter than 30 nucleotides were discarded after trimming with fastxtend tool available here https://github.com/warner-benjamin/fastxtend). In the last step, reads that were mapped to the Enterobacteria phage PhiX174 genome (GenBank: NC_001422.1) were discarded using bowtie2 v2.2.9 (-L 31-mp 4-rdg 6,6-local-no-unal) [24]. Remaining rRNA reads were removed using SortMeRNA v2.1 and SILVA databases version 119 [25,26].

### Transcriptome assembly

Healthy, exposed and infected readsets were co-assembled using Trinity v2.11.0 [27], then isoforms were clustered using SuperTranscripts [28]. Transcripts larger than 250 bases were retained. The taxonomic affiliation of assembled transcripts was performed with DIAMOND BLASTx (v2.1.8) against the marine eukaryote database MarFERRet v1.1 [29] and NCBI NR database v2025-02-12 [30]. For each transcript, a lowest common ancestor (LCA) was calculated with all matches with an e-value below 10^-5^ and a score above 95% of the best match (diamond option -outfmt 102 and -top 5). Functional annotation was conducted on six-frame translated sequences using multiple databases and tools: DIAMOND blastP against the KEGG protein database v2024-07-26, InterProScan [31] v5.61-93.0 including Gene Ontology (GO) term search, eggNOG mapper [32] v2.1.12 and its database v 5.0.2 [33], and the HMM search tool KofamKoala v1.3.0 to find KEGG Orthologues [34]. We used KEGG BRITE hierarchical classification systems [34] to attribute a global functional classification to each transcript. Functional and taxonomic information for each transcript are available in Table S1 and S2.

RNAseq reads were mapped back to the reference transcriptome using BWA-MEM2 v2.2.1 [35], then sorted with SAMtools [36]v1.18. Primary read alignments with more than 95% of identity over 80% of their length and a minimum of 100bp were selected. Read counts for each transcript were calculated with samtools idxstat. Using R v4.4.1, we selected transcripts assigned to the three dominant taxonomic groups: Bacillariophyta (mostly assigned to *Coscinodiscus* l *Bacillariophyta*), Oomycetes (mostly assigned to *L. coscinodisci)*, and *Pedinellales*. The Sankey plot was done with the R package network3D.

### Differential expression analyses

Differential expression analyses were performed using the DESeq2 R package [37]. Gene expression data were first filtered to retain only transcripts with sufficient abundance, defined as having at least 100 reads in one or more samples. Taxonomic assignments were based on Diamond LCA annotations, and genes were classified into major groups of interest, including *C. granii* (Bacillariophyta), *L. coscinodisci*-related oomycetes (Oomycota, Saprolegniaceae), and Pedinellales (Dictyochophyceae). For interspecies comparisons such as Pearson correlation analyses, genes were further filtered to include only those with a minimum coverage of 500 reads in at least one sample. Photosynthesis-related genes within the Pedinellales group were identified based on our annotation with specific Gene Ontology (GO) terms associated with photosynthetic processes.

BUSCO completeness was evaluated using **BUSCO v5.2.2** in transcriptome mode, with lineage-specific datasets: *stramenopiles_odb12* for *Coscinodiscus* and the Pedinellales, and *oomycota_odb12* for *L. coscinodisci*.

### Pedinellale identification from RNAseq reads

To taxonomically identify the Pedinellale protist present in the *C. granii* culture, we aligned RNAseq reads of the HEALTHY condition (replicate 1) on the full length 18S rRNA sequence of *Pteridomonas danica* (GenBank: PQ770944.1) and *C. granii* (AY485495.1) with BWA-mem2 (version 2.2.1). Reads aligned on P. *danica* 18S rRNA with more than 90% of identity over 80% of their length were retained, then a consensus sequence was builtd with bcftools consensus version 1.21 [38]. 242 complete 18S rRNAs sequences of Dictyochophytes available in the PR2 database [39] were aligned with mafft v7.49 using default options [40]. A maximum likelyhood phylogeny with 100 bootstrap was performed on MEGA version 11 [41].

### Structure prediction

The protein structure of the *C. granii* SSL protein was predicted using AlphaFold 3 [42]. The input amino acid sequence TRINITY_DN10640_c0 was used to generate a 3D structural model in PDB format.

## Results and discussion

### Construction of three transcriptomes in *Coscinodiscus/L. coscinodisci* cultures

We conducted a *de novo* co-assembly of the 9 RNA readsets (healthy, exposed, and infected conditions in triplicates)and obtained a metatranscriptome of 470,730 transcripts. These transcripts were subsequently clustered into 342,581 non-redondant super-transcripts larger than 250bp. We identified 15,712 transcripts affiliated to diatoms, including 7,595 with *Coscinodiscus* as best match and the others matching distant diatoms transcriptomes and genomes and therefore affiliated more broadly to Coscinodiscophyceae (2,203 transcripts) or Bacillariophyta (5,878 transcripts) (Table I, Figure 2). Similarly, 12,130 transcripts correspond to *L. coscinodisci* (Oomycota phylum) with 3,807 transcripts attributed to Oomycota and 4,542 to Saprolegniaceae. Finally, 9,250 transcripts were attributed to the Pedinellales (Dictyochophyceae), with 6,631 of them affiliated to *Pteridomonas danica*. An additional 5,045 eukaryotic transcripts could not be precisely assigned, likely due to incomplete reference transcriptomes or highly conserved proteins, identical between distant taxa. The other 300,444 transcripts did not match any known species and therefore could not be included in the following analyses

**Table I.** Summary of transcriptome assembly statistics and BUSCO completeness scores. The number of sequences, the N50 and the GC content of the transcripts assigned to each organism after de novo assembly are indicated. BUSCO shows the percentage of single-copy universal genes detected using the most appropriate lineage-specific dataset for each organism, namely *stramenopiles_odb12* for *Coscinodiscus* and the *Pedinellales*, and *oomycota_odb12* for *L. coscinodisci*.

| Organism | Transcripts | N50 | %GC | BUSCO (dataset) |
| --- | --- | --- | --- | --- |
| *C. granii* | 15,712 | 19,153 | 45.67% | 80.6% (stramenopiles_odb12) |
| *L. coscinodisci* | 12,130 | 11,344 | 32.87% | 66.2% (oomycota_odb12) |
| *Pedinellales* | 9,250 | 20,616 | 44.63% | 80.6% (stramenopiles_odb12) |

**Figure 2:**
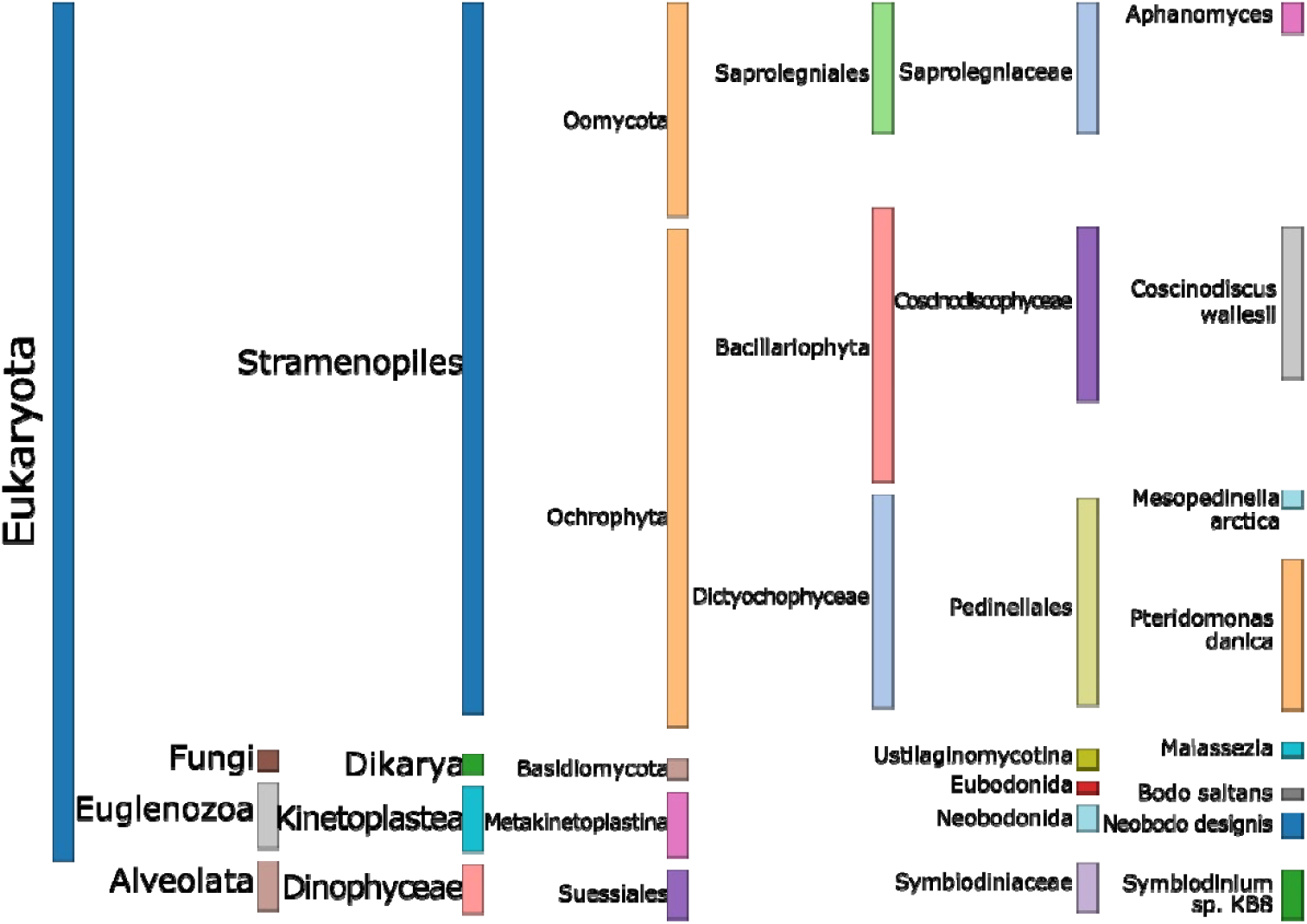
Taxonomic affiliation of eukaryote transcripts assembled from *C. granii* cultures. (Healthy, Exposed, and Infected). Only taxa represented by at least 500 genes are displayed.

**Figure 3:**
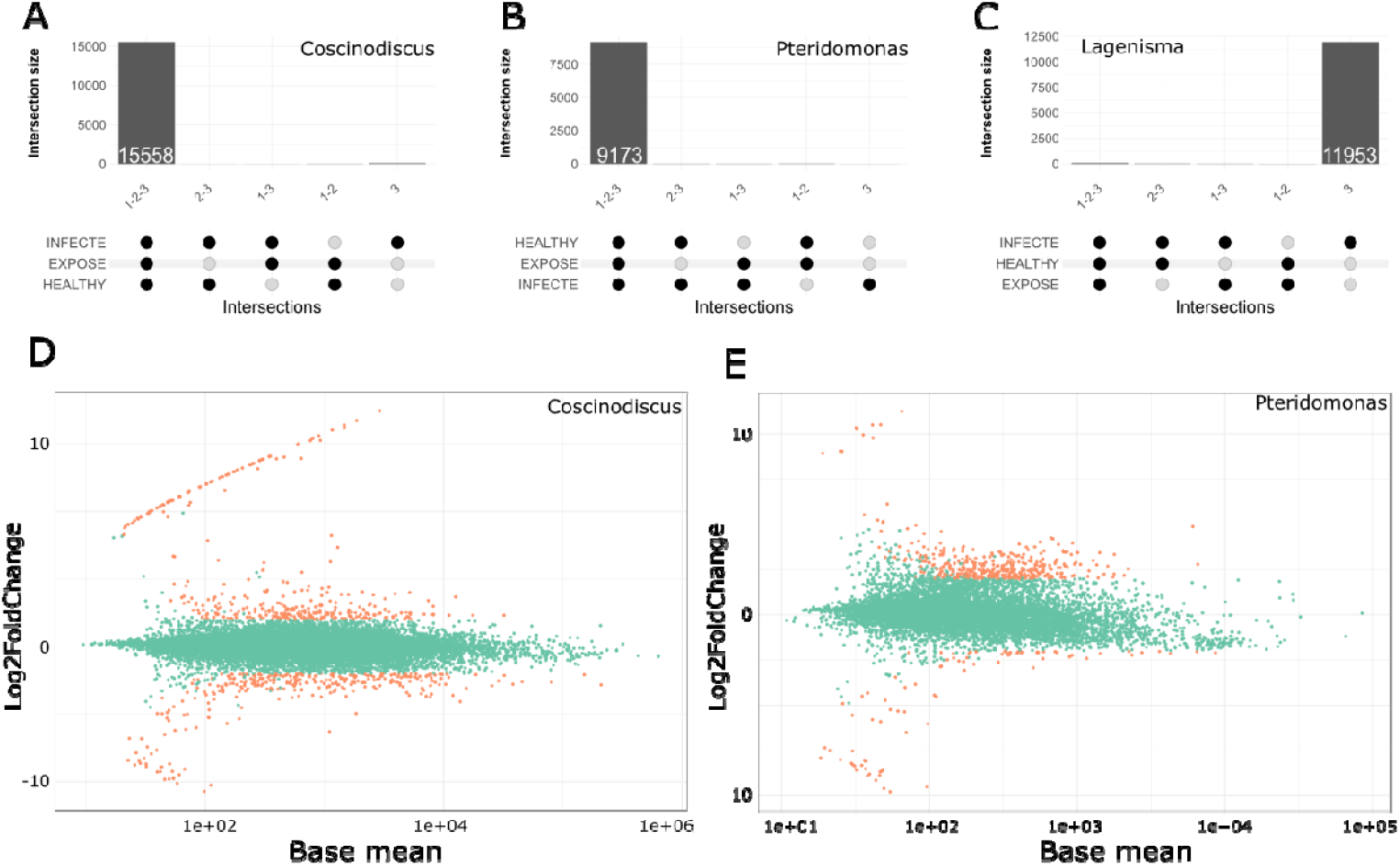
Gene expression of *C. granii, L. coscinodisci*. **(A–C)** UpSet plots showing the number of genes expressed in one or several conditions for *C. granii* (A), *Pteridomonas sp*. (B) and *L. coscinodisci* (C). Bar heights represent the size of each intersection, and dots below indicate the conditions where the genes are expressed. **(D–E)**: MA plots displaying log2 fold changes between healthy and infected cells versus the mean gene expression (log counts). Orange points represent significantly up- or down-regulated genes for *C. granii* (D) and *Pteridomonas sp*. (E).

To precise the taxonomy of the co-occuring pedinellales, we built an 18S rRNA consensus sequence from RNAseq reads and performed a phylogeny of all Dictyochophyceae 18S rRNA. The pedinellale sequenced in this experiment is closely related to *Pteridomonas danica* but with 3% of divergence on the 18S sequences. Therefore, we hereafter refer to it as *Pteridomonas* sp.. *P. danica* has lost photosynthesis genes [43] and possesses a reduced plastid genome [44]. Ecologically, *Pteridomonas sp*. function as important microbial consumers, preying on bacteria and other protists, thereby contributing to nutrient recycling and carbon cycling in marine ecosystems.

The *C. granii* and *Pteridomonas* sp. transcriptomes exhibited high completeness (80.6% complete BUSCOs) and low fragmentation (9% and 6.3% fragmented genes, respectively). In contrast, the *L. coscinodisci* transcriptome displayed lower completeness (66.2%) and higher fragmentation (12.7%). The lower BUSCO score observed for L. coscinodisci does not necessarily reflect an incomplete transcriptome due to many low-expressed genes, but rather a reduced genome resulting from the parasite’s reliance on host genes for a number of functions, as previously reported in some symbionts [45,46].

### Infection effectors identified in *L. coscinodisci* transcriptome

To address the repertoire of *L. coscinodisci* ‘weapons’ mobilized upon *C. granii* infection, we searched the L. coscinodisci transcriptome for sequences annotated as effector proteins typically associated with plant-infecting oomycetes. We have particularly focused on the Crinkler, cystatin, and RxLR families, as well as transposable element machinery, silencing machinery and cyclophilins (Table II, Table S3).

**Table II:**
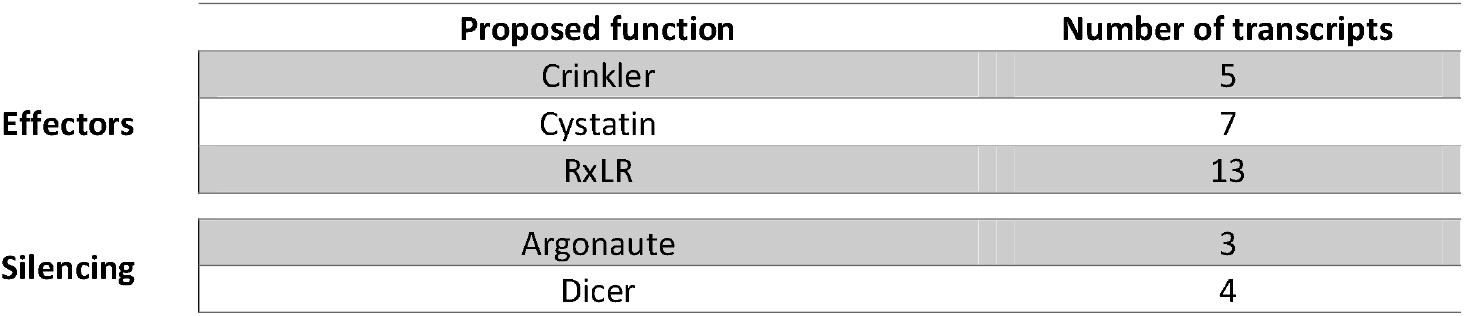
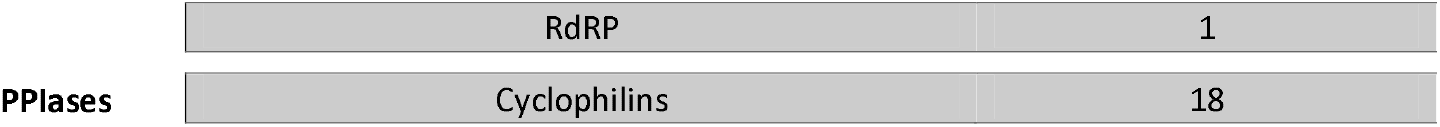
Number of oomycete transcripts assigned to putative functional categories. Classification was based on any matching functional annotation (best homologous hit in blastNR, KEGG, InterProScan, eggNOG, or KofamKoala, see material and method for details). Counts correspond to transcripts identified in infected samples. (PPIases: peptidyl-prolyl cis-trans isomerases, RdRP RNA-dependent RNA polymerase)

Crinkler effectors are secreted by oomycete pathogens, such as *Phytophthora* species, and play a crucial role in promoting infection by manipulating host plant cellular processes. Cinkler effectors like CRN12_997 from *Phytophthora capsici* target transcription factors such as SlTCP14-2 in tomatoes, inhibiting their DNA-binding ability and thus diminishing the plant’s immune response [47]. PsCRN108 from *Phytophthora sojae* can reprogram plant gene expression by targeting promoters of heat shock proteins, thereby suppressing stress responses and enhancing pathogen virulence [48]. Pi22798 (RxLR) from *Phytophthora infestans*, promotes the homodimerization of the host transcription factor StKNOX3, enhancing plant susceptibility by modulating transcriptional regulation [49]. HaRxL10 from *Hyaloperonospora arabidopsidis* targets the transcriptional repressor JAZ3 to activate jasmonic acid signalling and suppress salicylic acid-mediated defenses [50]. In *Hyaloperonospora arabidopsidis*, the oomycete RxLR effector HaRxL10 directly targets the central *Arabidopsis* circadian clock component CHE (CCA1 HIKING EXPEDITION), stabilizing it by inhibiting its degradation via the E3 ligase ZEITLUPE, which paradoxically suppresses CHE function, reprograms circadian gene expression, and enables the pathogen to repress plant immunity while manipulating growth and flowering processes [51].

Certain cyclophilins (CYPs) are implicated as virulence factors in oomycete infections as peptidyl-prolyl cis-trans isomerase (PPIase) [52]. This study identified 16 CYP orthogroups across 21 oomycete species, with each species encoding 15 to 35 CYP genes. Notably, three orthogroups contained proteins with signal peptides, indicating a potential role in secretion. Expression analysis in *Plasmopara infestans* and *Pl. halstedii* revealed distinct expression profiles during different asexual life stages, highlighting their importance in pathogenicity. We identify 18 cyclophilins transcripts in *L. coscinodisci* transcriptome.

SiRNA are described in plants oomycete as tuning the expression of oomycetes effectors and directly involved in virulence by targeting host mRNA. Among *L. coscinodisci* expressed genes, we could identify several key components of the RNA silencing machinery representing xx% of total gene expression. These genes include RNA-dependent RNA polymerases (RdRP, RDR2, RDR6), Argonaute1 (AGO1), a central component of the RNA-induced silencing complex (RISC), as well as argonaute linker 1 and 2 suggesting active silencing in *L. coscinodisci*. In addition, 2 genes have DUF1785 domain which is require in *A. thaliana* sRNA Duplex Unwinding [53].

Oomycete pathogens, such as *P. sojae*, secrete hundreds of effector proteins, including avirulence (Avr) proteins, that manipulate host immunity to promote infection. In a study by Wang et al 2019, it was discovered that endogenous small RNAs (sRNAs), derived from natural antisense transcripts, regulate the expression of the Avr1b gene in *P. sojae* [54]. The study demonstrated that sense and antisense expression of Avr1b is influenced by 10-base deletions in promoter regions, affecting the generation of these sRNAs. Further genome-wide analysis revealed that up to 31% of RXLR effector genes in *P. sojae* are associated with sRNAs. These sRNAs likely contribute to effector gene silencing and expression plasticity, suggesting a broader role in the pathogen’s ability to adapt and evade host immunity. Moreover, insertion/deletion variants of 9–10 base pairs were significantly enriched in the regulatory regions of RXLR genes, highlighting a mechanism by which bidirectional transcription and sRNA-mediated regulation shape effector expression patterns and, potentially, virulence.

In *A*.*thaliana*, the exchange of small RNAs (sRNAs) between the pathogen and the host, known as cross-kingdom RNA interference (ck-RNAi), leads to the silencing of host specific function [55]. For instance, the oomycete *Hyaloperonospora arabidopsidis* utilizes its sRNAs to bind to the plant’s Argonaute (AGO) proteins, part of the RNA-induced silencing complex, to suppress host immunity and promote virulence [55].

### Gene expression variations between healthy, exposed and infected cells

Global transcriptional patterns were compared across sample types using Pearson correlation of gene expression levels within each lineage. For both *C. granii* and *Pteridomonas sp*., healthy and exudate-exposed samples showed highly similar profiles (persons mean 0.97 and 0.93, respectively), indicating limited transcriptomic response in the absence of direct infection. Infected samples, however, displayed markedly distinct expression patterns (persons mean 0.9 and 0.76, respectively), consistent with expected major transcriptional reprogramming in the *L. coscinodisci* infected samples.

To determine the trophic mode of the co-occurrent *Pteridomonas sp*., we searched for photosynthetic genes in its transcriptome. GO term analysis (Table S3) revealed no genes associated with photosynthesis in this lineage, in stark contrast to *C. granii*, which shows a full set of photosynthesis-related GO terms including “photosynthesis” (GO:0015979), “photosynthetic electron transport” (GO:0019684), and “chlorophyll binding” (GO:0016168). Thus, the *Pteridomonas sp*. present in our culture is likely a heterotrophic or mixotrophic lineage that do not express photosynthetic genes in our conditions, consistent with known diversity of trophic strategies within the group.

Following *L. coscinodisci* infection, we identified 340 up-regulated genes and 304 down-regulated genes in *C. granii*, and 392 up-regulated and 87 down-regulated genes in *Pteridomonas sp*.. The differential gene expression analysis revealed no significant gene expression difference between healthy and exposed samples. Both *C. granii* and the *Pteridomonas sp*. appear unable to sense the cellular lysate of *C. granii* infected by *L. coscinodisci*.

### *Pteridomonas sp*. but not C. *granii* has a typical pathogen-exposition response

Eukaryotic cells have evolved sophisticated signalling mechanisms to respond to various biotic stresses, including infections by oomycete pathogens. In plants, a key signalling pathway involved in this response is the Mitogen-Activated Protein Kinase (MAPK) cascade [56]. In contrast to *C. granii*, the MAPK signalling pathway is activated in *Pteridomonas sp*. with 16 genes having a log2 fold change (log2fc) up to 3.7. These findings could suggest that *Pteridomonas sp*. detects the parasite and mount a MAPK-mediated defense response.

Treatment of plant cells with a *Phytophthora sojae* protein elicitor induces a biphasic cytosolic Ca^2+^ increase, which coincides with a burst of reactive oxygen species (ROS) and phytoalexin accumulation [57]. Calcium-dependent protein kinases (CDPKs) then transduce the signal to defense effectors. In our study, a strong calcium signalling response is observed in *Pteridomonas sp*. exposed to *L. coscinodisci*, with 13 calcium-related genes upregulated (log2fc up to 3.7). In contrast, *Coscinodiscus* shows a minimal response, with only one gene upregulated (log2fc 2.17). In potatoes, StCDPK7 is upregulated upon *P. infestans* infection and phosphorylates phenylalanine ammonia -lyase in a Ca^2+^-dependent manner [57]. Similarly, in pepper, the Ca^2+^ sensor CBL1 and its interacting partner CIPK1 are essential for resistance to *P. capsici. A* loss of CaCIPK1 increases susceptibility, while its overexpression enhances defense responses, including ROS production and cell death [57].

Interestingly, our transcriptomic data reveal that 16 genes associated with exosome-related functions (as annotated by BRITE category C) are upregulated in *Pteridomonas sp*., and six related genes are upregulated in *C. granii*.

The role of extracellular vesicles (EVs) in plant immunity has gained growing interest in recent years. For decades, plant EVs have been observed to accumulate in infected cells, particularly beneath sites of pathogen attack and at papillae formation zones, where they are believed to contribute to the establishment of defensive barriers [58]. EVs have also been detected surrounding fungal haustoria and within the extrahaustorial matrix. These vesicles, which may carry antimicrobial compounds or RNAs, are hypothesized to deliver their contents into fungal haustoria either through membrane fusion or endocytic uptake. This raises the question of whether EVs also exist in algae and what roles they might play. It is plausible that algal EVs carry defense-related molecules such as enzymes or small RNAs. Interestingly, Deng et al. (2024) showed that the diatom *Coscinodiscus radiatus* produces EVs during recovery from nutrient stress, which help eliminate harmful metabolites and support cell rejuvenation. Notably, this vesicle production is triggered by bacterial signals, suggesting a broader role for EVs in stress responses and microbial interactions [59].

In *Pteridomonas sp*., a substantial transcriptional reprogramming of motility-related genes is observed, with strong induction (up to +4.6-fold change) of 47 cytoskeleton-associated genes and 93 cilium-related genes, including members of the dynein and kinesin families. This shift in gene expression likely reflects one or both of two non-mutually exclusive strategies: an evasive behavioural response triggered by chemical cues released following parasite-induced cell lysis and/or a broader adaptation of motility and signalling pathways in response to environmental changes. Notably, the absence of significant upregulation in nutrient transporter genes suggests that this response is not driven by opportunistic feeding behaviour but aligns more closely with an active escape mechanism in reaction to infection-associated exudates. In parallel, several mitochondrial genes also exhibit strong upregulation, with log2 fold changes ranging from +2.2 to +4.9, indicating a possible increase in energy demand. This mitochondrial activation may support enhanced motility and cellular remodelling.

### Genome stress, reparation, and ribonucleases in C. granii

*C. granii* does not exhibit this early infection-related signalling response despite showing a phenotype of infected cells. Instead, *C. granii* upregulates a variety of ribonucleases, notably members of the Ribonuclease H (RNase H) superfamily, with eight proteins showing fold changes up to 16.7. These enzymes specifically target RNA/DNA hybrid structures, indicating genomic stress. The response also includes upregulating four DNA repair genes, with log2fc ranging from 2.3 to 3.7 (Figure 4 and Table S3).

**Figure 4:**
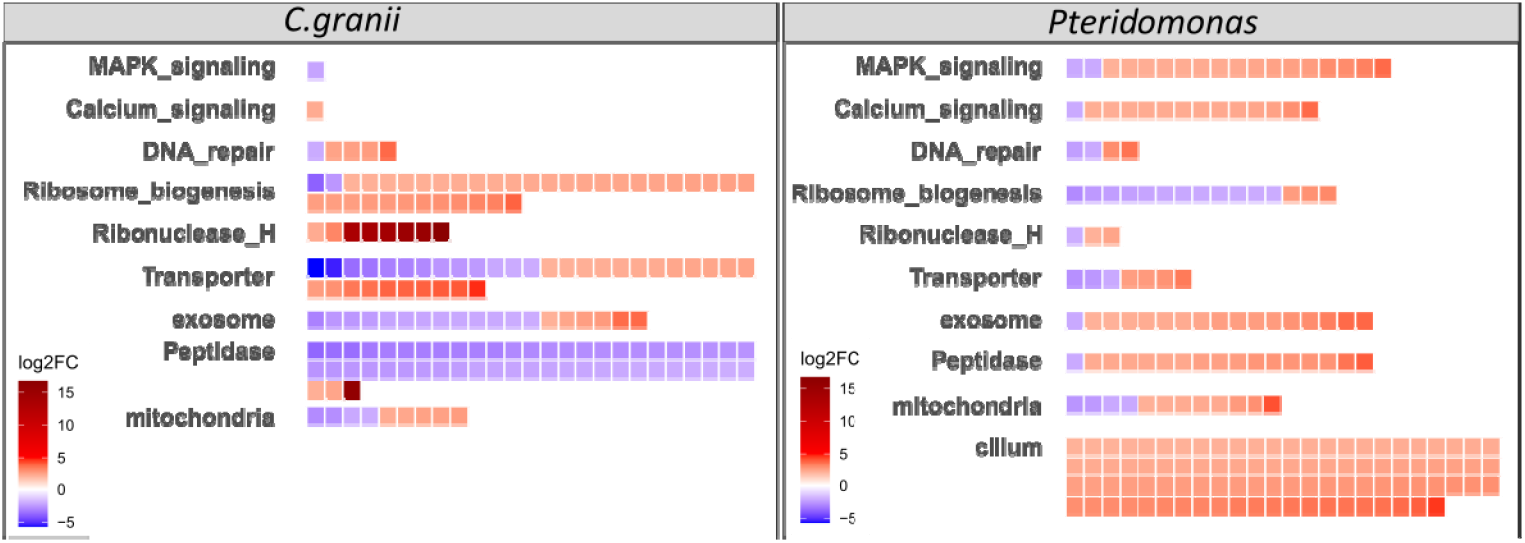
Overexpressed functions in *C. granii* and *Pteridomonas sp*. in *C. granii*-infected samples. Each coloured box represents a differentially regulated gene, classified according to its functional annotation using InterPro (IPR) domains (*Ribonuclease H, Peptidase*) or KEGG Brite category (Brite C) (*Transporter [BR:ko02000], DNA repair [BR:ko03400], Ribosome biogenesis [BR:ko03009], Peptidase [BR:ko01002], MAPK signaling [BR:ko04010], Calcium signaling [BR:ko04020], Cilium [BR:ko04961], Exosome [BR:ko04147], Mitochondria [BR:ko00190]*)

This observation raises the intriguing possibility that *C. granii* may contribute to host defense by degrading foreign RNA hybridized to regulatory regions of host DNA. R-loops are structures where RNA anneals to one strand of DNA leaving the other strand single-stranded, they are known to induce genomic stress and require timely resolution to maintain genome [60]. Additionally, ribosome biogenesis in *C. granii* appears significantly upregulated during oomycete infection, with 35 genes upregulated (max lox2fc 3.85). This observation mirrors findings in plants, where ribosome biogenesis is a crucial cellular process frequently targeted by oomycete effectors. For example, the *Phytophthora infestans* effector Pi23226 localizes to the plant nucleolus and binds the 3⍰ end of 25S rRNA precursors, disrupting pre-rRNA processing, inducing nucleolar stress, and impairing global translation [61].

The ribosomal upregulation observed in *C. granii* is likely a compensatory mechanism in response to parasite-mediated inhibition of ribosome function, possibly involving parasite-derived cyclophilins that could interfere with ribosomal activity and protein synthesis.

### C.granii peptidase shifts during infection

We could identify 460 peptidases-encoding genes in the *C. granii* transcriptome, that is consistent with the 461 peptidase genes found in *P. tricornutum* [62]. 32 peptidases are downregulated and 3 of them upregulated. Proteases are known to play key roles in numerous cellular processes, including protein turnover and stress response pathways. Notably, eight of these proteases are also classified as molecular chaperones, suggesting that the observed response is more likely indicative of a broad physiological shift linked to cellular stress rather than a specific reaction to pathogen attack [63].

In contrast, a Papain-like cysteine peptidase is one of the most upregulated proteins during infection (log2fc 16). Papain-like cysteine peptidases are crucial in plant-pathogen interactions, notably during oomycete infections [64]. These enzymes are part of a broader family of cysteine proteases involved in plant defense mechanisms by degrading pathogen proteins and activating immune responses. Further, a study on the parasite *Amoebophrya* revealed that dinoflagellates can produce lytic compounds as a defensive strategy [65]. However, the precise structure and class of these compounds remained unidentified. This raises the possibility that proteases could be among these elusive defensive agents. In the *L. coscinodisci* transcriptome, we could identify 7 cystatin-like proteins that could represent potential candidates to counter *Coscinodiscus* Ppapain-like cysteine peptidases. Indeed, the production of cystatin-like protease inhibitors in several pathogens has been shown to ? counteract these defenses. *Phytophthora palmivora* secretes PpalEPIC8, a cystatin-like inhibitor that targets papain in papaya, thereby enhancing the pathogen’s virulence by inhibiting the plant’s defense protease activity [66]. Similarly, *P. infestans*, responsible for late blight in tomatoes and potatoes, produces EPIC2B, which inhibits a novel papain-like protease, PIP1, involved in plant defense [67]. In *Arabidopsis*, the XYLEM CYSTEINE PEPTIDASE 1 (XCP1) and its inhibitor CYSTATIN 6 (CYS6) regulate pattern-triggered immunity by modulating the stability of the NADPH oxidase RBOHD, which is crucial for reactive oxygen species (ROS) production during immune responses [68].

### β-carbolines biosynthesis putative enzyme

β-carbolines alkaloids have been previously detected by mass spectrometry in algal cells infected with *L. coscinodisci* [69] suggesting a defensce function during the infection. In plants, β-carboline alkaloids are synthesised by the Strictosidine synthase enzyme, which catalyses the condensation of secologanin and tryptophan to form strictosidine [70,71]. Interestingly, strictosidine synthase-like (SSL) enzymes in *A. thaliana* have also been implicated in biotic stress responses [72]. SSL presents a stable architecture composed of six radially arranged β-sheets.

By generating protein structure of unannotated or DUFs proteins in our dataset *in-silico* we identified as a six-bladed β-propeller fold one of the most expressed gene (5^th^ by basemean) and strongly upregulated protein (log2fc = 8.2) annotated as DUF839 (TRINITY_DN10640_c0_g2). Thus, it is conceivable that this DUF839-domain protein may participate in the β-carboline biosynthetic pathway. However, this hypothesis remains speculative and requires further biochemical and functional validation because this fold, conserved across kingdoms, supports a broad spectrum of enzymatic activities, including catalysis, molecular recognition, and signal transduction [73].

## Conclusion

The *C*.*granii/L. coscinodisci* pathosystem presents a highly promising model for studying marine oomycete-diatoms interactions, particularly due to its notable parallels with well-characterized plant–pathogen interactions. These transcriptomes are a first reference to understand molecular mechanisms supporting *L. coscinodisci* infection and *C. granii* response. We observed a rich genetic toolkit that enables a sophisticated infection strategy resembling those of plant-infecting oomycetes, including hyphal development and the deployment of conserved molecular effectors such as Crinklers, cystatins, RxLR proteins, and transposable elements. Additionally, it encodes components of the RNA interference machinery: Argonaute, Dicer, and RdRP as well as cyclophilins.

The activation of an exosome-mediated response to infection or pathogen-associated elicitors suggests an ancient defense mechanism, potentially predating and informing those found in terrestrial plants. This intriguing possibility warrants further investigation.

Moreover, this pathosystem opens new avenues for characterizing enigmatic protein families, such as the SSL-like proteins and DUF839, not only in algae but also within broader lineages.

In conclusion, this model stands out for its relative simplicity compared to land plant systems while still encompassing key aspects of host-parasite interactions, particularly within the context of exceptionally large diatom cells, which can span hundreds of microns. Sequencing genomes for the host (*C. granii*) and the parasite (*L. coscinodisci*) will be crucial to unlocking the system’s full potential. Ultimately, this marine microalga model offers a unique and powerful platform to investigate fundamental questions in evolutionary parasitology, effector biology, and host manipulation.

## Acknowledgments

We acknowledge the Max Planck Institute for Chemical Ecology for hosting the research where the strains are permanently maintained and Ute Kieb for providing Helgoland natural seawater every month since 8 years.

We acknowledge the staff of the sequencing platform at Genoscope for their attentiveness and their efforts in seeking technological solutions to experimental challenges and the Scientific Information Technology Laboratory (LIS) for support in high-performance computing at Genoscope.

## Data availability

Raw sequencing reads have been deposited in the ENA (EMBL-EBI) under accession number PRJEB102303. Assembled transcriptomes are available at https://doi.org/10.57745/X0W739.

